# Selective Regulation of Tuft Cell-Like Small Cell Lung Cancer by Novel Transcriptional Co-activators C11orf53 and COLCA2

**DOI:** 10.1101/2022.07.12.494433

**Authors:** Chen Zhou, Hui Huang, Yunyi Wang, Erdem Sendinc, Yang Shi

**Author notes:** Correspondence: Yang Shi. These authors contributed equally: Hui Huang, Yunyi Wang.

## Abstract

The DepMap project has generated a huge resource for investigating selectively essential genes, which represent potential cancer therapeutic targets. However, manually sorting out which of the hundreds of selectively essential genes is understudied and warrants investigations is time-consuming and probably not practical. To efficiently identify uncharacterized, selectively essential genes, we collected and ranked the 347 selectively essential genes from the DepMap dataset by their PubMed publication counts, based on the assumption that genes with low publication counts are un-studied or under-studied. We successfully validated two of the top candidates in our ranking system, C11orf53 and COLCA2, as new vulnerabilities that are selectively essential in the class II POU domain transcription factor POU2F3-dependent tuft cell-like small cell lung cancer (SCLC) cell lines. Importantly, we found that the sequence motif, which mediates physical interactions of the transcriptional co-activator POU2AF1 with two of the class II POU domain-containing family of transcription factors (POU2F1 and POU2F2), is also present in the N-terminal regions of C11orf53 and COLCA2. We further confirmed that 1) COLCA2 physically interacts with POU2F3 through this conserved sequence motif; 2) this interaction is important for COLCA2 to regulate tuft cell-like SCLC cell growth, and 3) both C11orf53 and COLCA2 contain transcriptional co-activator domains. Consistently, we find similar transcriptomic changes in response to the loss of COLCA2 or POU2F3 in SCLC cells. In summary, our analysis pipeline enables identification and prioritization of understudied but important, selectively essential genes, leading to the identification of two new transcriptional co-activators for the class II POU domain transcription factors. Disruption of this important physical interaction is predicted to be a potential therapeutic strategy to selectively inhibit tuft cell-like SCLCs.

---

Dear Editor,

An ideal cancer therapy drug should kill the cancer cells while displaying limited and controllable toxicities towards normal cells^1^. Therefore, genes that are selectively essential in cancer but not normal cells are good therapeutic targets. However, while genetic dependencies in many different cancer cell lines have been defined through genome-wide genetic screens, much less was known about the genetic dependency for normal cells of different tissue origins at the genome-wide level. To overcome this problem, the DepMap project has developed a method to identify genes that are selectively essential only in a subset of cancer cell lines but not others by using the genome-wide CRISPR-based fitness screen data from ∼1000 human cancer cell lines^2–4^. DepMap calculates a CERES score for each gene in every cancer cell line, which measures the effect of CRISPR-mediated gene knock-out on cell fitness. A CERES score of 0 indicates that a gene is non-essential in a given cell line and a score of -1 is the median of all common essential genes. To identify selectively essential genes, DepMap calculates a NormLRT (Normality Likelihood Ratio Test) score for each gene by using its CERES scores across different cancer cell lines. A higher NormLRT score indicates that the distribution of CERES scores of this gene across different cancer cell lines is more deviated from normal distribution, suggesting that the gene may be more selectively essential. If a gene is not essential in a large number of cancer cell lines originated from different tissue types, it is less likely that it’s a part of a core pathway critical for normal cells. Identification and characterization of these selectively essential genes will not only deepen our understanding of cancer biology, but also guide cancer drug development. However, manually sorting out which of the selectively essential genes is understudied and warrants investigations is time-consuming and probably not practical.

To prioritize understudied selectively essential genes, we first recalculated the NormLRT score for each gene, since the NormLRT scores are not directly available from the DepMap portal. By using a threshold (NormLRT > 125) reported previously^4^, we collected 347 potential selectively essential genes. We then ranked them according to the PubMed publication count^5^. A gene with a low PubMed publication count is often understudied, and in our ranking system, the top 9 most understudied genes are C11orf53, C3orf38, TMEM164, ZNF511, KCNK13, BEST3, CYB561A3, C12orf49 and COLCA2 (Fig. 1a). In fact, CYB561A3 has only recently been identified as the key lysosomal iron reductase and a novel cancer vulnerability in Burkitt lymphoma^6^. This demonstrated that our strategy can efficiently identify novel potential cancer therapeutic targets.

**Fig. 1.**
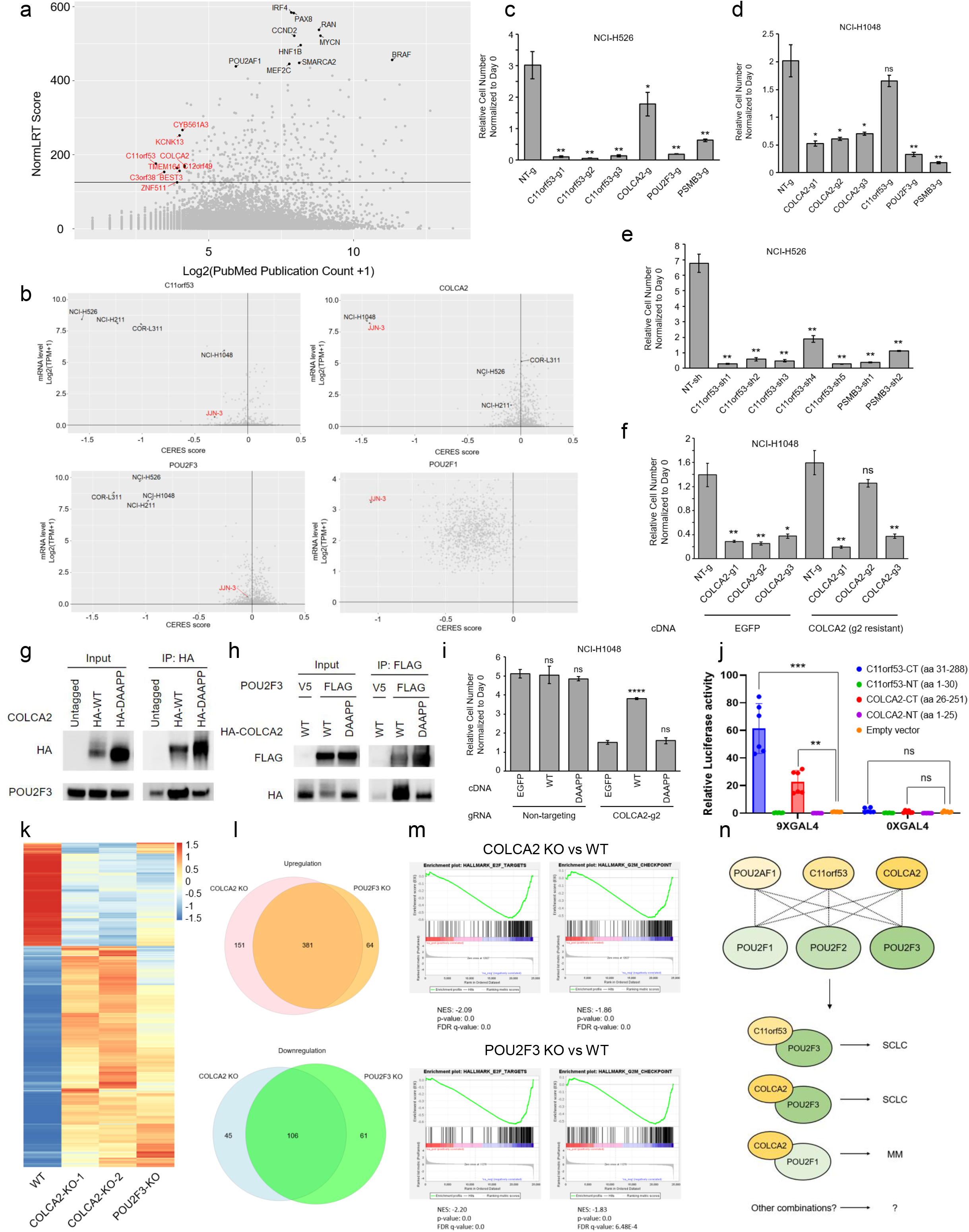
Selective regulation of tuft cell-like small cell lung cancer by novel transcriptional co-activators C11orf53 and COLCA2. **a** Scatter plot showing each gene’s NormLRT score vs log2(PubMed publication count+1). Dots labeled with text in black are the top 10 genes with the highest NormLRT scores. Dots labeled with text in red are the top 9 selectively essential genes with the lowest PubMed publication counts. The black horizontal line is NormLRT=125 threshold. **b** Scatter plots showing the expression levels of C11orf53/COLCA2/POU2F3/POU2F1 in different cancer cell lines vs the corresponding CERES scores. The plots were generated by using DepMap 21Q2 dataset. SCLC cell lines are labeled with text in black. JJN-3, a multiple myeloma cell line, is labeled with text in red. **c** Cell-Titer Glo assay result of EGFP-expressing NCI-H526 cells infected with different Cas9-sgRNA lentiviruses. EGFP-expressing NCI-H526 cells were infected with Cas9-sgRNA lentiviruses, selected with puromycin and seeded in 96-well plate (same number of cells). Cell-Titer Glo assay was performed on day 0 and day 5. sgRNA targeting PSMB3, a common essential gene, was a positive control. NT: non-targeting. n=3, two-tailed unpaired Student’s t test. **P < 0.01; *P < 0.05. **d** Cell-Titer Glo assay result of NCI-H1048 cells infected with different Cas9-sgRNA lentiviruses. NCI-H1048 cells were infected with Cas9-sgRNA lentiviruses, selected with puromycin and seeded in 96-well plate (same number of cells). Cell-Titer Glo assay was performed on day 0 and day 4. sgRNA targeting PSMB3, a common essential gene, was a positive control. NT: non-targeting. n=3, two-tailed unpaired Student’s t test. **P < 0.01; *P < 0.05. **e** Cell-Titer Glo assay result of NCI-H526 cells infected with different shRNA lentiviruses. NCI-H526 cells were infected with shRNA lentiviruses and 2 days later the same number of cells were seeded in 96-well plate. Cell-Titer Glo assay was performed on day 0 and day 4. shRNAs targeting PSMB3, a common essential gene, served as positive control. NT: non-targeting. n=3, two-tailed unpaired Student’s t test. **P < 0.01. **f** Cell-Titer Glo assay result of EGFP- or COLCA2 (resistant to COLCA2-gRNA-2)-expressing NCI-H1048 cells infected with different Cas9-sgRNA lentiviruses. EGFP- or COLCA2 (resistant to COLCA2-gRNA-2)-expressing NCI-H1048 cells were generated first. They were then infected with Cas9-sgRNA lentiviruses, selected with puromycin and seeded in 96-well plate (same number of cells). Cell-Titer Glo assay was performed on day 0 and day. NT: non-targeting. n=3, two-tailed unpaired Student’s t test. **P < 0.01; *P < 0.05. Statistical analysis was done within each cDNA group. **g** Immunoprecipitation-western blot result for exogenously expressed HA-COLCA2 (WT or mutant where VKELL is mutated to DAAPP) and POU2F3 in HEK293T cells. **h** Immunoprecipitation-western blot result for exogenously expressed FLAG-POU2F3 and HA-COLCA2 (WT or mutant where VKELL is mutated to DAAPP) in HEK293T cells. **i** Cell-Titer Glo assay result of EGFP- or COLCA2 (resistant to COLCA2-gRNA2)-WT/mutant (VKELL mutated to DAAPP)-expressing NCI-H1048 cells infected with different Cas9-sgRNA lentiviruses. EGFP- or COLCA2 (resistant to COLCA2-gRNA2)-WT/mutant (VKELL mutated to DAAPP)-expressing NCI-H1048 cells were generated first. They were then infected with Cas9-sgRNA lentiviruses, selected with puromycin and seeded in 96-well plate (same number of cells). Cell-Titer Glo assay was performed on day 0 and day 5. NT: non-targeting. n=3, two-tailed unpaired Student’s t test. ****P < 0.0001. Statistical analysis was done within each gRNA group. **j** Luciferase reporter assay where the N-terminal (NT) and C-terminal (CT) domains of C11orf53/COLCA2 were fused to GAL4 DNA binding domain and the luciferase reporter gene was driven by a minimal promoter downstream of 9xGAL4 binding sites or 0xGAL4 binding sites. n=6, two-tailed unpaired Student’s t test. ***P < 0.001; **P < 0.01. **k** Heatmap showing normalized expression levels of significantly up- and down-regulated genes in COLCA2 KO vs WT NCI-H1048 cells in different samples. **l** Venn diagrams showing the overlap of up- or down-regulated genes in COLCA2 KO and POU2F3 KO NCI-H1048 cells. **m** GSEA analysis results for gene expression changes in COLCA2 or POU2F3 KO NCI-H1048 cells vs WT cells. **n** The model, in which any of POU2AF1/C11orf53/COLCA2 might interact with any of POU2F1/2/3. Different complexes may be formed in different contexts to execute their biological functions.

We subsequently focused on C11orf53 and COLCA2, which stand out as attractive candidates because of the significant CERES scores in their dependent cell lines where they are highly expressed, suggesting strong growth phenotypes (Fig. 1b). Remarkably, guided by the top co-dependencies pre-calculated by DepMap, we found that among the ∼1,000 cancer cell lines, C11orf53 is only essential in the DNA-binding transcription factor POU2F3-dependent small cell lung cancer (SCLC) cell lines and one of the only two COLCA2-dependent cell lines is also a POU2F3-dependent SCLC cell line. Specifically, C11orf53 is essential for the SCLC cell lines, COR-L311, NCI-H526 and NCI-H211 while COLCA2 is essential in NCI-H1048. The reason for this apparent, mutually exclusive requirement is likely due to their differential expression. In the C11orf53-dependent SCLC cell lines, C11orf53 expression is higher than that of COLCA2, while in the COLCA2-dependent SCLC cell lines, COLCA2 expression is higher (Fig. 1b). In addition to C11orf53 and COLCA2, POU2F3 is selectively essential in all four of these SCLC lines. The POU2F3-dependent SCLC was reported previously and it was characterized as a tuft cell-like variant SCLC^7^. However, the mechanistic details of how POU2F3 drives expression of tuft cell-specific gene expression programs remains unclear. Based on our analysis, we hypothesized that C11orf53/COLCA2 and POU2F3 may work in the same pathway to regulate growth of a subset of SCLC, which are tuft cell-like.

To confirm the DepMap data, we knocked out C11orf53 and COLCA2 in NCI-H526 cells and NCI-H1048 cells by CRISPR using different sgRNAs, respectively, and performed cell growth assays. Consistent with the DepMap data, three different C11orf53 gRNAs significantly decreased NCI-H526 cell growth (Fig. 1c). The COLCA2 gRNA also decreased NCI-H526 cell growth but to a lesser extent than the C11orf53 gRNAs. Similarly, three different COLCA2 gRNAs, but not the C11orf53 gRNA, significantly decreased the growth of NCI-H1048 cells (Fig. 1d). Cas9 has been shown to cause cell toxicity when generating DNA double stranded breaks at loci with high copy numbers^8,9^. Since there is an increase in copy number at the C11orf53 locus in NCI-H526 cells (Supplementary Fig. S1), we wished to rule out the possibility that the cell growth defect is caused by the generation of double stranded breaks by Cas9. Consistent with the CRISPR approach, five different C11orf53 shRNAs also caused growth defects of NCI-H526 cells (Fig. 1e), supporting the notion that it’s the loss of C11orf53 that caused the cell growth phenotype. Similar results were also obtained in COR-L311 cells (Supplementary Fig. S2). We also carried out genetic rescue experiments in the COLCA2 gRNA-treated NCI-H1048 cells, whose growth phenotype was rescued by a gRNA-resistant COLCA2 transgene encoding wildtype COLCA2 protein (Fig. 1f). These findings suggest that C11orf53 and COLCA2 are required for SCLC cell growth.

We next investigated the molecular functions of C11orf53 and COLCA2 by performing protein domain analysis through the Pfam portal^10^ (Supplementary Fig. S3). Interestingly, sequence alignment analysis revealed that both C11orf53 and COLCA2 contain an N-terminal sequence, (R/K)xYQGVRVKxxVK(D/E)LLxx(K/R)R, which is also present in the transcriptional co-activator POU2AF1 (Supplementary Fig. S4). Importantly, this motif has been shown to mediate physical interactions of POU2AF1 with the highly-conserved POU-specific domains (Supplementary Fig. S5) found in the POU domain class 2 family of DNA-binding transcription factors, POU2F1 and POU2F2^11,12^. As discussed, POU2F3, which is the third member of this family, is essential for all four of the tuft cell-like SCLC cell lines showing C11orf53- or COLCA2-dependency in DepMap (Fig. 1b), leading to our hypothesis that C11orf53 and COLCA2 may act as co-activators of POU2F3 and regulate transcription of genes critical for growth and survival of a subset of SCLC represented by these four SCLC lines.

Our co-activator hypothesis predicts that C11orf53 and COLCA2 physically interact with POU2F3. Indeed, co-immunoprecipitation experiments in HEK293T cells successfully detected physical interactions between COLCA2 and POU2F3 (Fig. 1g, h). Furthermore, mutations in the predicted interaction motif of COLCA2 resulted in a reduced interaction with POU2F3, supporting our model that COLCA2 and POU2F3 interact with each other through the conserved motif (Fig. 1g, h). Importantly, the mutant COLCA2 with a reduced ability to physically interact with POU2F3 also showed compromised ability to rescue the growth defect in NCI-H1048 cells caused by COLCA2 loss, indicating physical interaction between COLCA2 and POU2F3 is important for SCLC cell growth regulation (Fig. 1i). As co-activators, COLCA2 and C11or53 are also predicted to carry transcriptional activation domains. Indeed, when the C-terminal region of COLCA2 (aa 26-251) or C11or53 (aa 31-288) is fused to the DNA-binding domain of GAL4, the fusion proteins activated transcription of the luciferase reporter gene in a GAL4 binding sites-dependent manner (Fig. 1j). Collectively, these findings identify COLCA2 (and likely C11orf53) as a co-activator for POU2F3 and demonstrate their physical interaction is critical for the growth of NCI-H1048 cells.

To explore the molecular basis underlying the growth defect of COLCA2-deficient NCI-H1048 cells, we performed RNA-sequencing. Consistent with the co-activator hypothesis, most of the genes with significant changes in the COLCA2-deficient cells (adjusted p-value < 0.05, 1208 upregulated genes, 563 downregulated genes) also showed similar changes in the POU2F3-deficient cells (Fig. 1k). In addition, there was a large overlap between the up- or down-regulated genes (adjusted p-value < 0.05, fold change > 2) in the COLCA2-deficient cells and those in the POU2F3-deficient cells (Fig. 1l). POU2F3 was reported to drive expression of tuft cell markers^7^. Consistently, loss of COLCA2 also decreased the expression of tuft cell markers (Supplementary Fig. S6). Importantly, GSEA analysis found that the downregulated genes in the COLCA2-deficient and POU2F3-deficient cells were significantly enriched in cell cycle related pathways (Fig. 1m), many of which are known positive regulators of cell cycle, including CDC25A, CENPE and KIF15 (Supplementary Fig. S7), which may explain the growth phenotype associated with the loss of COLCA2 or POU2F3. These observations provide further functional evidence supporting the model that COLCA2 functions as a co-activator for POU2F3.

In summary, using the PubMed publication count we were able to prioritize understudied potential cancer therapeutic targets, leading to the identification of C11orf53 and COLCA2 as two of the top candidates in our ranking system. We subsequently validated both genes as novel vulnerabilities in a subset of SCLC cell lines that are tuft-cell like. We provided further biochemical and functional data demonstrating that COLCA2 functions as a co-activator for POU2F3 to drive transcriptional program important for growth of tuft cell-like SCLC. The highly selective nature of these co-activators in cancer coupled with the previous reports that both Colca2^-/-^ mice and C11orf53^-/-^ mice are viable^13,14^ suggest that disrupting the interactions of these coactivators with POU2F family of transcription factors could be a viable therapeutic strategy with minimal toxicities. While our work was ongoing, two papers, one in bioRxiv^15^ and the other in Nature^14^ appeared online where the authors also investigated the role of C11orf53 (renamed as POU2AF1)^15^ and both C11orf53 (renamed as POU2AF1/OCA-T1) and COLCA2 (renamed as POU2AF2/OCA-T2)^14^ in SCLC, and our conclusion is essentially the same as those reached by these investigators. We noticed that in addition to SCLC cells, COLCA2 is also essential in a multiple myeloma cell line, JJN-3, where POU2F1 instead of POU3F3, is highly expressed and essential (Fig. 1b). This suggests that in JJN-3 cells, COLCA2 may work as a co-activator for POU2F1 to promote cancer cell growth. Based on the findings discussed above, we propose a general model (Fig. 1n). In this model, POU2AF1/C11orf53/COLCA2 are a family of co-activators for the POU domain transcription factors, POU2F1/2/3, to regulate expression of genes critical for growth of a subset of SCLCs, and possibly additional cancer such as multiple myeloma. How the co-activators are paired with these DNA-binding transcription factors is likely dictated by the relative expression levels of these factors. We further noticed that C11orf53 and COLCA2 are expressed in multiple tumor samples across different cancer types, and their levels of expression are much higher than those in normal tissues in a subset of patients (Supplementary Fig. S8), suggesting these two genes could have roles in cancers beyond tuft cell-like SCLC and multiple myeloma.

## Acknowledgements

This project is supported by the Ludwig Institute for Cancer Research. Y.S. is an American Cancer Society Research Professor.

## Author contributions

C.Z. and Y.S. conceived the project and wrote the manuscript with input from all the co-authors. C.Z. performed most of the experiments and did bioinformatic analysis. H.H. helped with molecular cloning and performed the dual-luciferase reporter assay. Y.W. performed the immunoprecipitation experiments. E.S. helped with the CellTiter-Glo assay.

---

**Supplementary Fig. S1.**
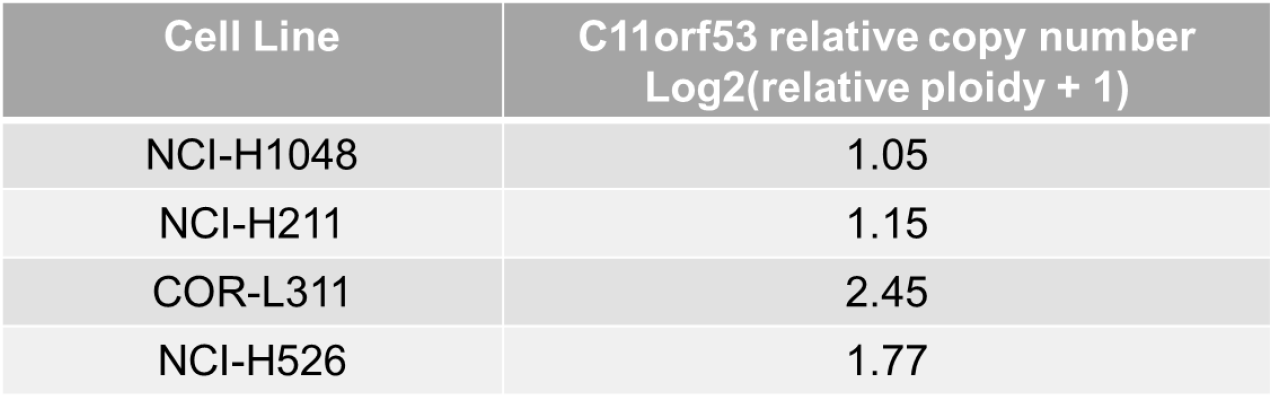
C11orf53 relative copy number in tuft cell-like SCLC cell lines. The copy number data was from DepMap 21Q2 dataset. Relative copy number = 1 means the copy number of this gene is the same as most of the rest of the genome.

**Supplementary Fig. S2.**
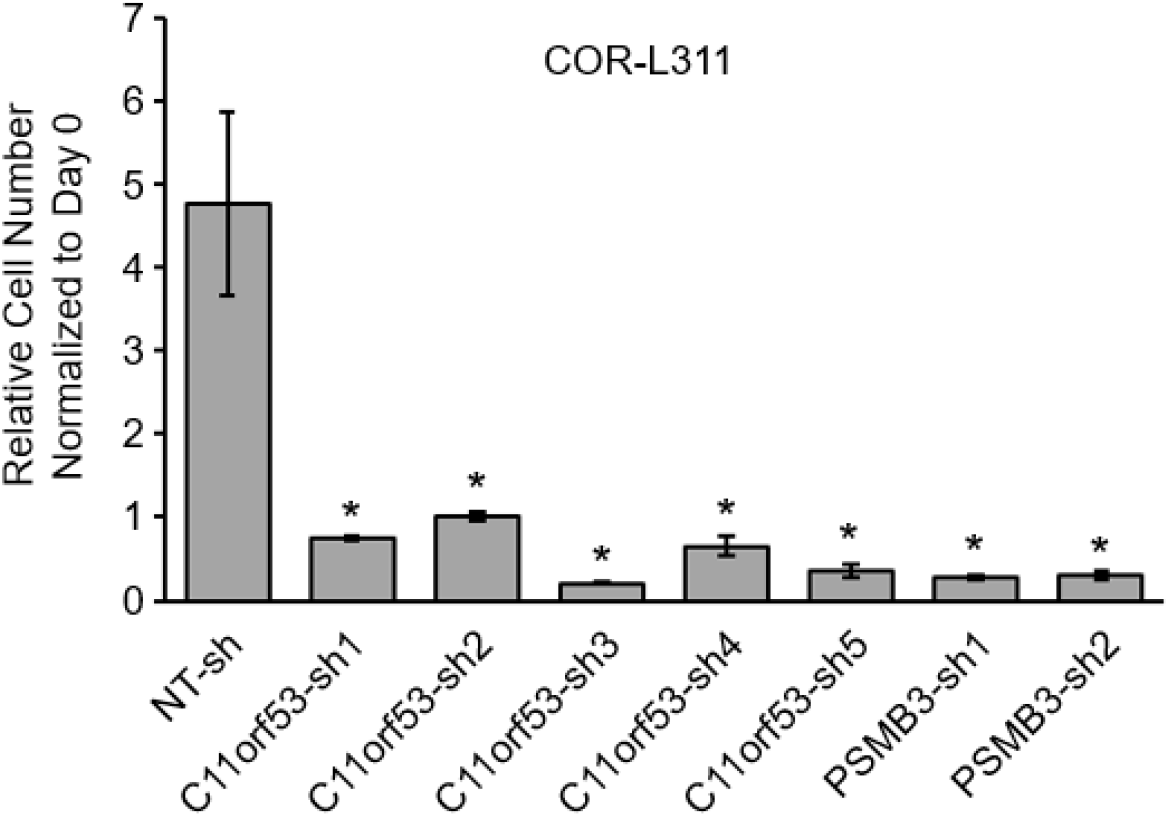
C11orf53 knock-down decreased growth of COR-L311 cells. Cell-Titer Glo assay result of COR-L311 cells infected with different shRNA lentiviruses. COR-L311 cells were infected with shRNA lentiviruses and 2 days later the same number of cells were seeded in 96-well plate. Cell-Titer Glo assay was performed on day 0 and day. NT: non-targeting. n=3, two-tailed unpaired Student’s t test. *P < 0.05.

**Supplementary Fig. S3.**
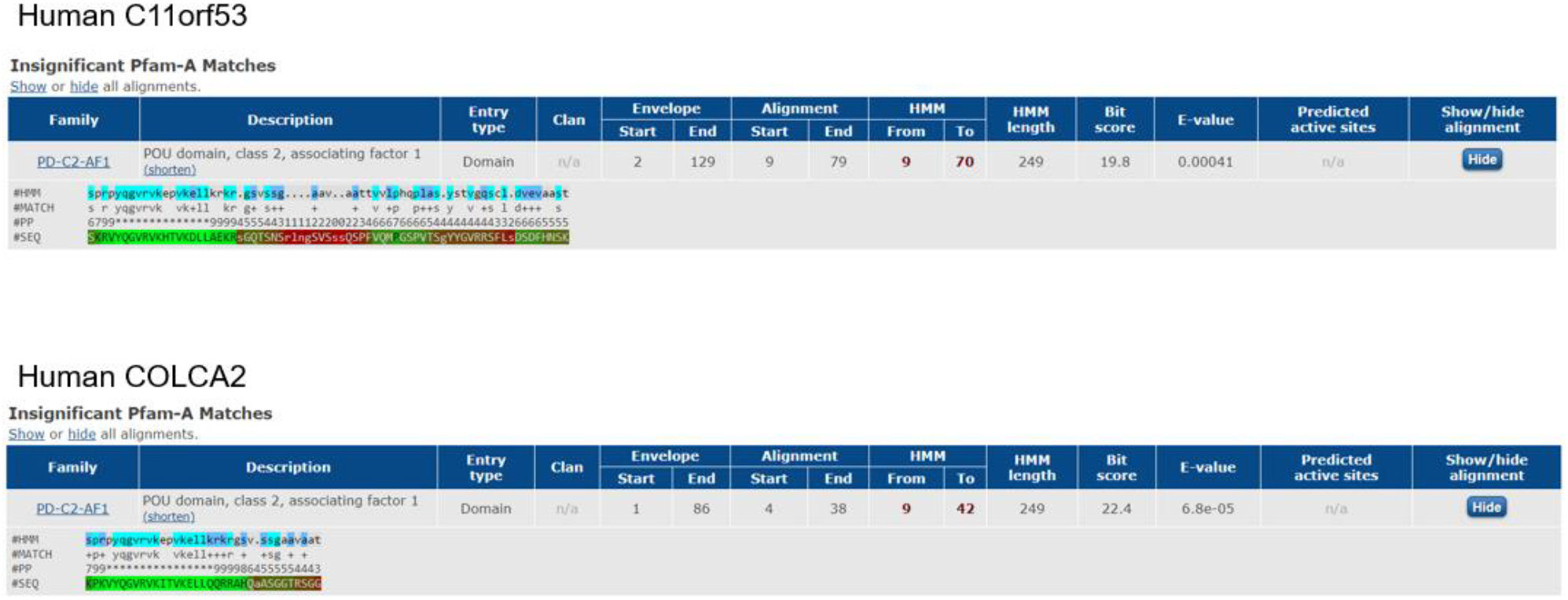
Pfam protein domain prediction result. Pfam was used to predict protein domains in human C11orf53 and COLCA2.

**Supplementary Fig. S4.**
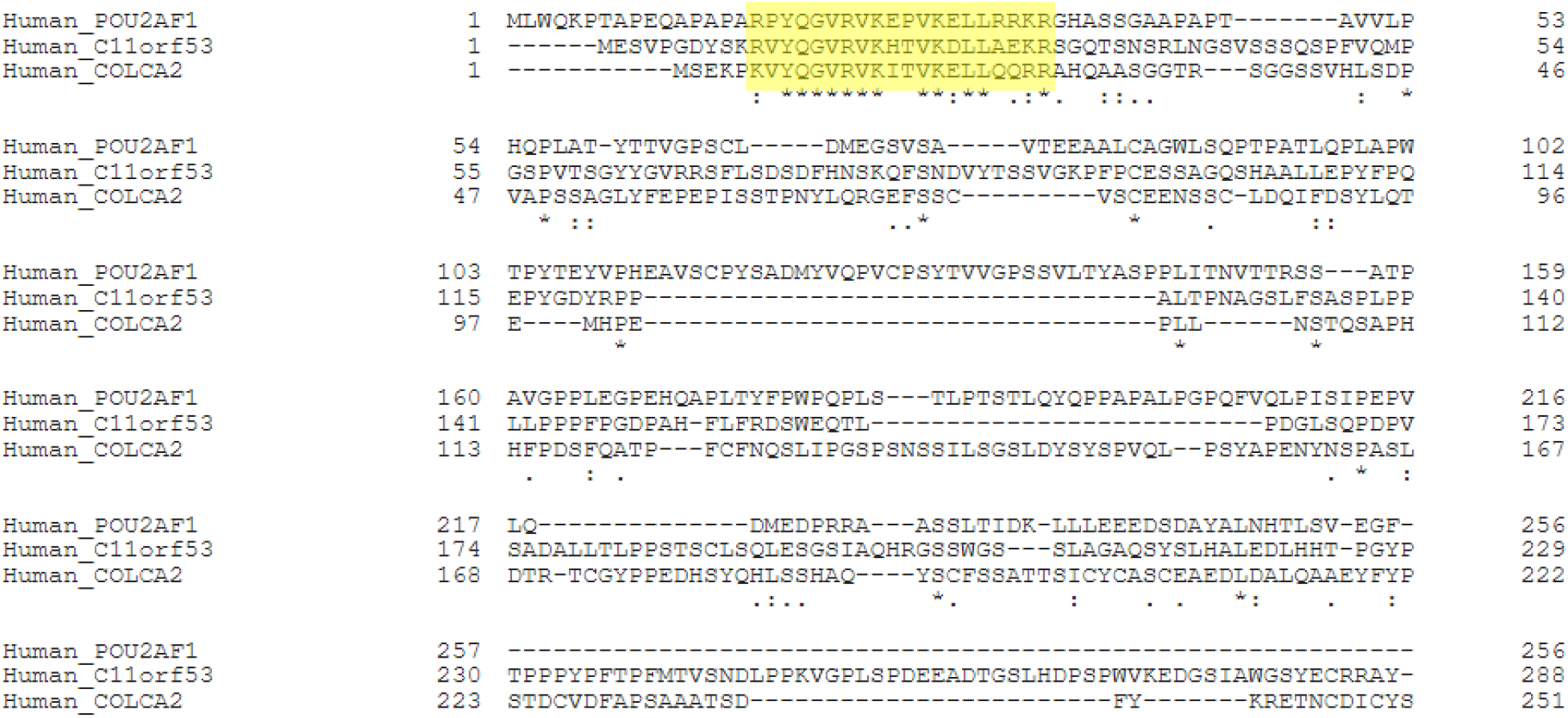
Protein sequence alignment result of human POU2AF1/C11orf53/COLCA2. Protein sequence alignment identifies a shared, conserved sequence motif (highlighted) at the N-terminal regions of human POU2AF1, C11orf53 and COLCA2.

**Supplementary Fig. S5.**
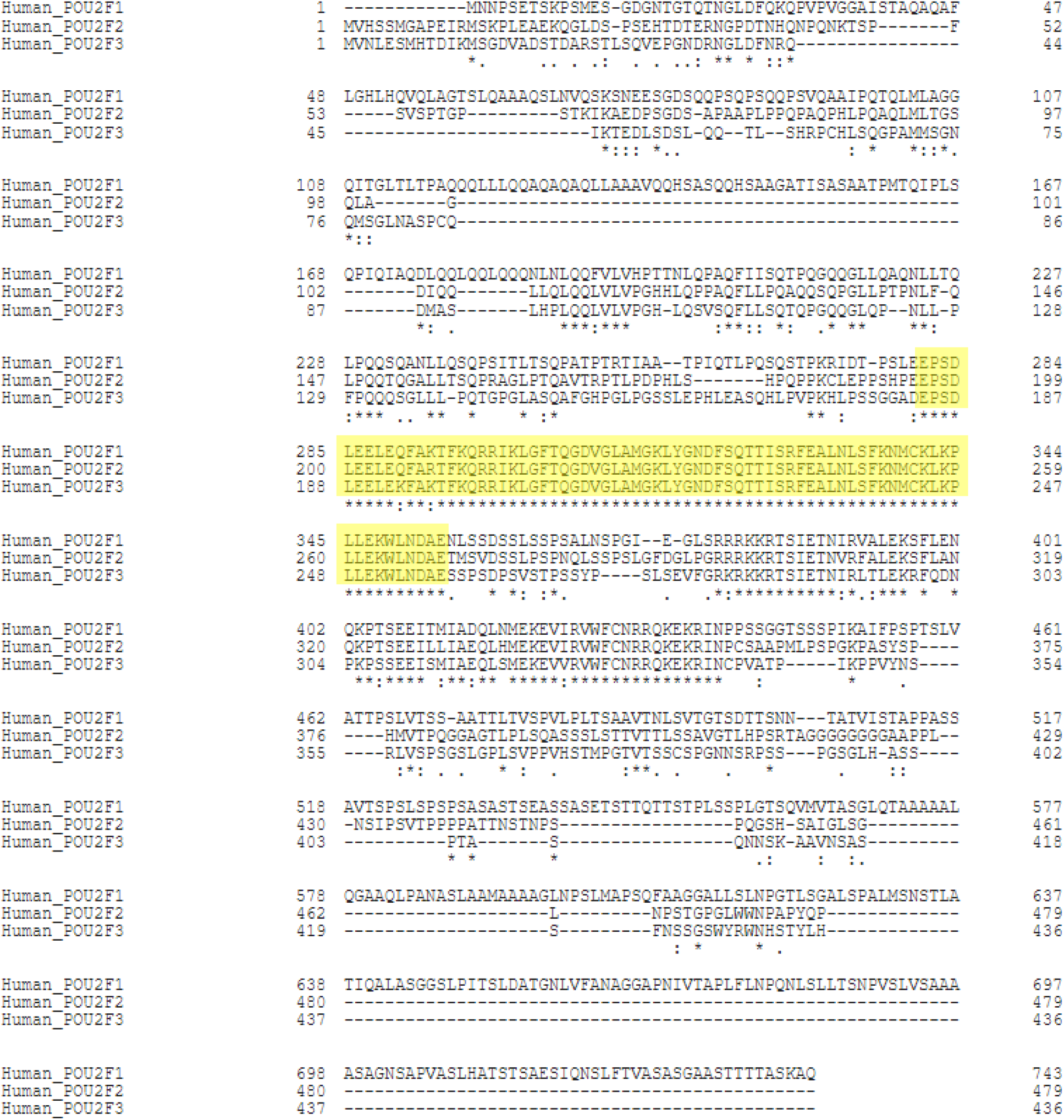
Protein sequence alignment result of human POU2F1/POU2F2/POU2F3. Alignment was performed using Uniprot. The conserved POU-specific domain is highlighted.

**Supplementary Fig. S6.**
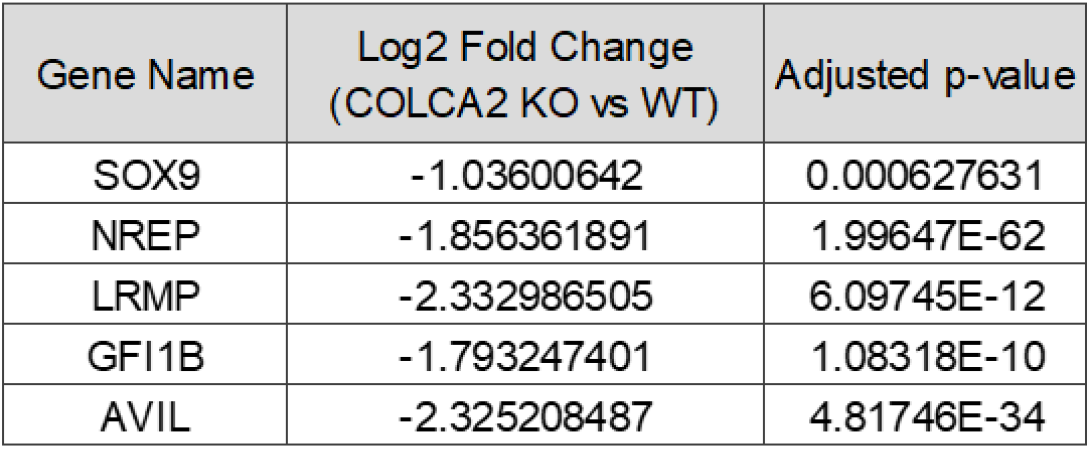
Gene expression changes of tuft cell markers in COLCA2 KO vs WT NCI-H1048 cells.

**Supplementary Fig. S7.**
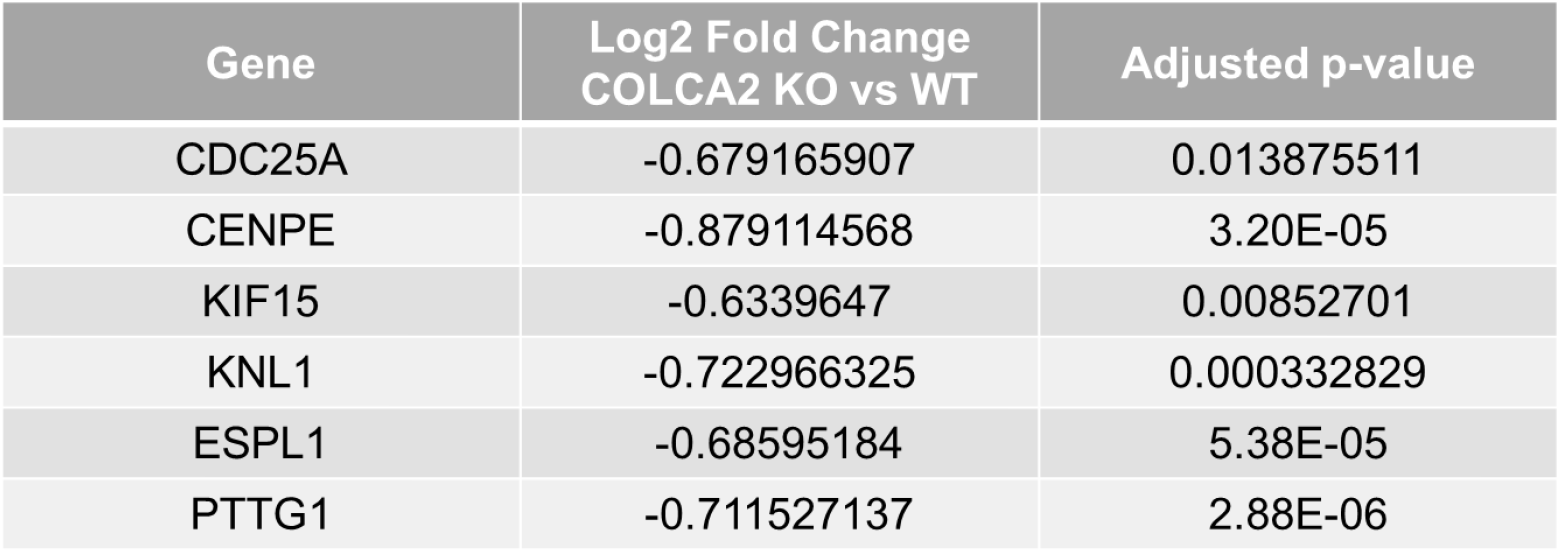
Gene expression changes of cell cycle regulators in COLCA2 KO vs WT NCI-H1048 cells.

**Supplementary Fig. S8.**
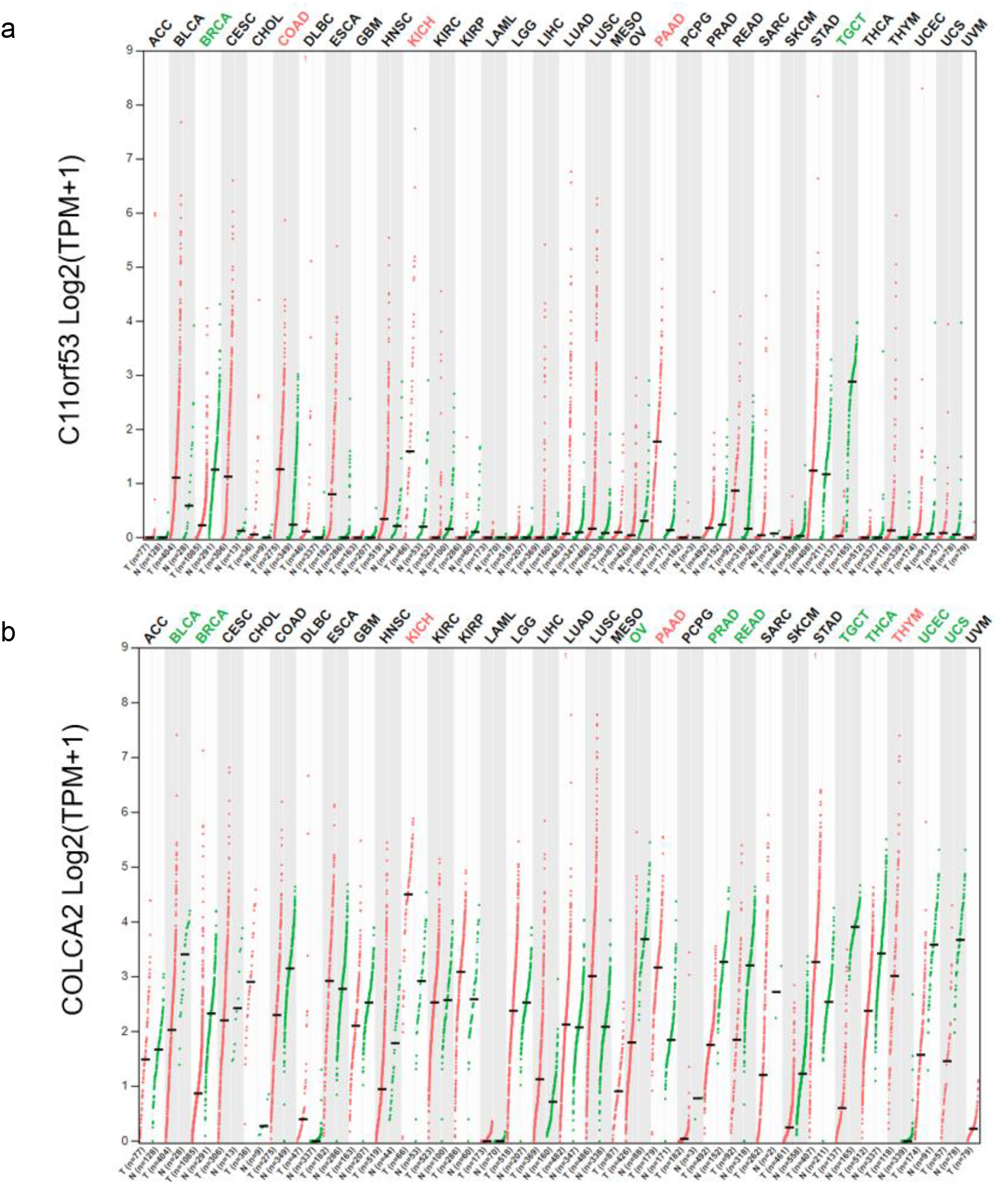
Gene expression level of C11orf53/COLCA2 in different tumor and normal samples. **a** Data from GEPIA showing that C11orf53 is highly expressed in a subset of tumor samples across different cancers. **b** Data from GEPIA showing that COLCA2 is highly expressed in a subset of tumor samples across different cancers. Red dots are tumor samples and green dots are normal samples.

## Supplementary Methods

### Cell lines

HEK293T, NCI-H1048 and NCI-H526 cells were obtained from ATCC. COR-L311 cells were obtained from ECACC. HEK293T cells were cultured in DMEM supplemented with 10% FBS and 1% penicillin/streptomycin. NCI-H1048 cells were cultured in DMEM/F12 supplemented with 5% FBS, 0.005 mg/mL insulin, 0.01 mg/mL transferrin, 30 nM sodium selenite, 10 nM hydrocortisone and 10 nM beta-estradiol. NCI-H526 cells and COR-L311 cells were cultured in RPMI1640 supplemented with 10% FBS. All the cells were cultured in a 5% CO2 incubator at 37°C.

### Antibodies

Antibodies used in this study include HA (Cell Signaling Technology #14031S, 1:1000 for western blot), FLAG (Sigma Aldrich #F1804, 1:1000 for western blot) and POU2F3 (Sigma Aldrich #HPA019562, 1:500 for western blot).

### Gene depletion by CRISPR/Cas9

The gRNA oligos, with sequences for their respective target genes listed below, were annealed and cloned into a lentiCRISPR v2 lentiviral vector. lentiCRISPR v2 was a gift from Feng Zhang (Addgene #52961)^1^. Lentivirus carrying lentiCRISPR v2 plasmid was produced by co-transfecting HEK293T cells with two packaging plasmids, psPAX2 (Addgene #12260) and pMD2.G (Addgene #12259, both were gifts from Didier Trono), and by harvesting viral supernatant after 48 h by passing through a 0.45 um filter. Collected lentivirus was used directly to infect NCI-H1048, NCI-H526 and COR-L311 cells with the addition of 4 ug/ml polybrene by centrifugation at 550 g for 30 min. 48h later, the infected cells were selected with puromycin for 4-5 days before being used for subsequent assays.

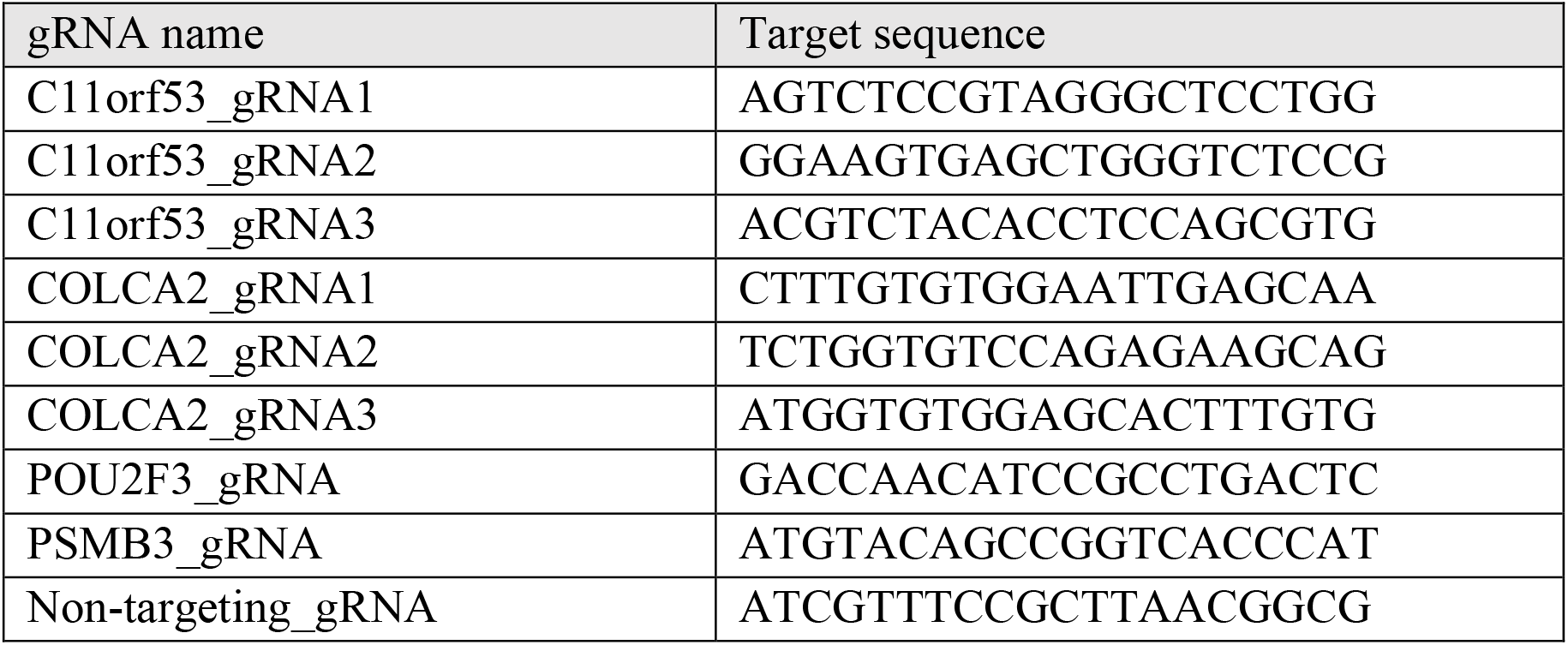

### Expression plasmid construction

COLCA2-gRNA2-resistant COLCA2 cDNA was amplified from cDNA library reverse transcribed from RNA extracted from NCI-H1048 cells. C11orf53 cDNA was amplified from cDNA library reverse transcribed from RNA extracted from NCI-H526 cells. EGFP cDNA, C11orf53 cDNA, untagged/HA-tagged WT/mutant (VKELL mutated to DAAPP) COLCA2-gRNA2-resistant COLCA2 cDNA were cloned into pCMV-blasticidin lentiviral vector by using NEBuilder HiFi DNA assembly. LentiV-Neo-POU2F3 was a gift from Christopher Vakoc (Addgene #122175)^2^. V5- or FLAG-tagged POU2F3 were cloned into pCMV-puromycin lentiviral vector by using NEBuilder HiFi DNA assembly.

### Stable cell line generation

Lentivirus derived from pCMV-blasticidin plasmid expressing WT or mutant COLCA2 and EGFP was produced by co-transfecting HEK293T cells with two packaging plasmids (psPAX2 and pMD2.G) and by harvesting viral supernatant after 48 h by passing through a 0.45 um filter. Collected lentivirus was used directly to infect NCI-H1048 cells with the addition of 4 ug/ml polybrene by centrifugation at 550 g for 30 min. 48h later, the infected cells were selected with 4 ug/mL blasticidin for 4-5 days before being used for subsequent assays.

### Gene knockdown by shRNA

The shRNA oligos, with their target sequences listed below, were annealed and cloned into a pLKO.1-Puromycin lentiviral vector (Addgene #10878, a gift from David Root)^3^. Lentivirus carrying pLKO.1 plasmid was produced by co-transfecting HEK293T cells with two packaging plasmids (psPAX2 and pMD2.G) and by harvesting viral supernatant after 48 h by passing through a 0.45 um filter. Collected lentivirus was used directly to infect NCI-H526 and COR-L311 cells with the addition of 4 ug/ml polybrene by centrifugation at 550 g for 30 min. 48h later, the infected cells were used for subsequent assays.

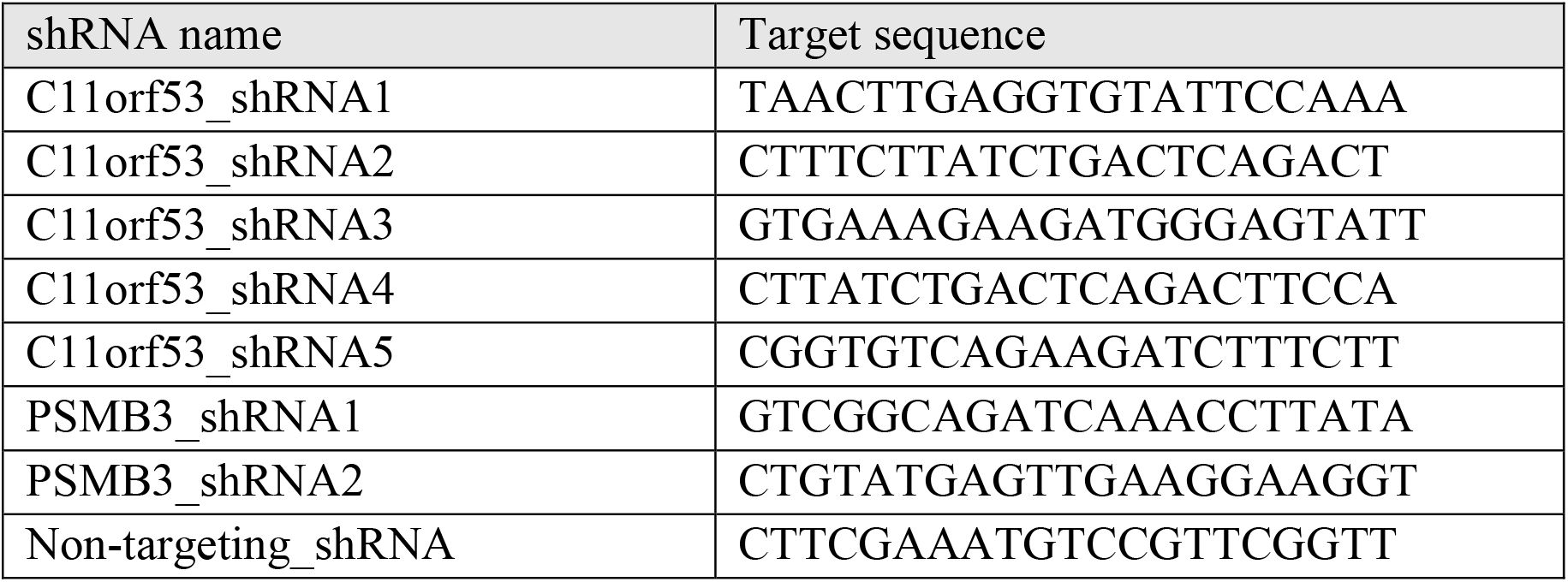

### Cell growth assay

6000 NCI-H1048/NCI-H526/COR-L311 cells were seeded in each well of 96-well plate for cell growth assay in 100 uL volume. 4-5 days later, relative cell number was assessed by using the CellTiter-Glo luminescent assay kit (Promega #G7572) according to the manufacturer’s instructions.

### Immunoprecipitation (IP)

HEK293T cells in a 10 cm dish were transfected with the indicated plasmids (10 ug each), and harvested 48 h post-transfection. Cell pellets were lysed for 30 min on ice in IP lysis buffer (20 mM Tris-HCl pH8.0, 150 mM NaCl, 1% Triton X-100, 1.5 mM MgCl_2_) supplemented with Complete EDTA-free protease inhibitor cocktail (Sigma-Aldrich, cat#5892791001) and PhosSTOP (Sigma-Aldrich, cat#04906837001). Protein lysates were cleared by 10 min centrifugation to pellet cell debris. Cleared protein lysates were incubated with anti-HA magnetic beads (Thermo Fisher Scientific, cat#88837) or anti-FLAG magnetic beads (Thermo Fisher Scientific, cat#A36798) overnight at 4 °C. Immunoprecipitants were then washed three times with the IP lysis buffer and eluted using 100 uL 2x SDS loading buffer.

### GAL4 dual-luciferase reporter assay

cDNA fragments encoding the C-terminal and N-terminal domains of C11orf53 and COLCA2 were cloned into the pFN26A (Cat#E1380, Promega) vector by Gibson Assembly (Cat#E2611L, NEB). C11orf53 N-terminal: aa 1-30. C11orf53 C-terminal: aa 31-288. COLCA2 N-terminal: aa 1-25. COLCA2 C-terminal: aa 26-251. The plasmids were then co-transfected with the pGL4.35[luc2P/9XGAL4 UAS/Hygro] vector (Cat#E1370, Promega) or the version without GAL4 binding sites into HEK293T cells. Specifically, 50 ng reporter plasmids was co-transfected with 100 ng pGL4.35 vector/the version without GAL4 binding sites into 10,000 HEK293T cells in each well of 96-well plate by lipofectamine 2000 (Cat#11668019, Invitrogen). Luciferase activity was measured by dual-luciferase reporter assay kit (Cat#E1910, Promega) 48 hours after transfection following the manufacturer’s instructions.

### RNA-seq and data analysis

Total RNA was extracted by using TRIzol/chloroform extraction method and further treated with DNase by using RNA Clean & Concentrator (Zymo Research #R1014). Poly(A)+ RNA was selected by using the NEBNext® Poly(A) mRNA Magnetic Isolation Module (NEB #E7490S). RNA-seq libraries were prepared according to the manual for NEBNext® Ultra™ II RNA Library Prep Kit (NEB #E7770L) for Illumina and sequenced on a NextSeq 2000 set at 50 PE mode. Read counts for genes were determined by using Salmon and differential gene expression analysis was done by using DESeq2.

### Identification of selectively essential genes

CRISPR gene effect table containing CERES scores was downloaded from DepMap Public 21Q2 dataset. To calculate NormLRT score^4^ for each gene, the distribution of CERES scores across all cell lines were fitted with both a normal distribution and a skew normal distribution. NormLRT score = 2*(log likelihood ratio of the fitted skew normal distribution – log likelihood ratio of the fitted normal distribution). PubMed publication count was derived from gene2pubmed file available on NCBI FTP.

